# SemEHR: A General-purpose Semantic Search System to Surface Semantic Data from Clinical Notes for Tailored Care, Trial Recruitment and Clinical Research

**DOI:** 10.1101/235622

**Authors:** Honghan Wu, Giulia Toti, Katherine I. Morley, Zina M. Ibrahim, Amos Folarin, Richard Jackson, Ismail Kartoglu, Asha Agrawal, Clive Stringer, Darren Gale, Genevieve Gorrell, Angus Roberts, Matthew Broadbent, Robert Stewart, Richard JB Dobson

## Abstract

**Objective:** Unlocking the data contained within both structured and unstructured components of Electronic Health Records (EHRs) has the potential to provide a step change in data available forsecondary research use, generation of actionable medical insights, hospital management and trial recruitment. To achieve this, we implemented SemEHR - a semantic search and analytics, open source tool for EHRs.

**Methods:** SemEHR implements a generic information extraction (IE) and retrieval infrastructure by identifying contextualised mentions of a wide range of biomedical concepts within EHRs. Natural Language Processing (NLP) annotations are further assembled at patient level and extended with EHR-specific knowledge to generate a *timeline* for each patient. The semantic data is serviced via ontology-based search and analytics interfaces.

**Results:** SemEHR has been deployed to a number of UK hospitals including the Clinical Record Interactive Search (CRIS), an anonymised replica of the EHR of the UK South London and Maudsley (SLaM) NHS Foundation Trust, one of Europes largest providers of mental health services. In two CRIS-based studies, SemEHR achieved 93% (Hepatitis C case) and 99% (HIV case) F-Measure results in identifying true positive patients. At King’s College Hospital in London, as part of the CogStack programme (github.com/cogstack), SemEHR is being used to recruit patients into the UK Dept of Health 100k Genome Project (genomicsengland.co.uk). The validation study suggests that the tool can validate previously recruited cases and is very fast in searching phenotypes - time for recruitment criteria checking reduced from days to minutes. Validated on an open intensive care EHR data - MIMICIII, the vital signs extracted by SemEHR can achieve around 97% accuracy.

**Conclusion:** Results from the multiple case studies demonstrate SemEHR’s efficiency - weeks or months of work can be done within hours or minutes in some cases. SemEHR provides a more comprehensive view of a patient, bringing in more and unexpected insight compared to study-oriented bespoke information extraction systems.

SemEHR is open source available at https://github.com/CogStack/SemEHR.

## BACKGROUND

The opportunity for secondary use of the wealth of information contained within Electronic Health Records (EHRs) has attracted researchers interested in investigating approaches to provide more tailored and timely care, improve efficiency of services and to derive new scientific and medical insights [1–4]. In addition to structured data contained within relational database tables (such as ICD-10 diagnoses codes), EHR documents are filled with unstructured clinical notes, such as nursing records, radiology reports or discharge summaries. These notes add a richness and depth to EHR-based studies [5–7], providing data and insight beyond what is available within the thin supernatant of data stored within the structured fields.

Deriving actionable insights from the EHR, including the unstructured component is challenging.It requires bringing together expertise in the clinical domain, the underlying healthcare information systems and text analytics techniques, e.g. Natural Language Processing (NLP). For example, the Clinical Record Interactive Search system (CRIS) [8], an anonymised replica of the EHR used in the South London and Maudsley NHS Foundation Trust (SLaM) in the UK, was designed for supporting clinical and scientific studies. Since its launch in 2009, a large number of studies ([9–13] to name a few) used the CRIS resource in conjunction with NLP or text mining techniques. Although these studies answered different clinical questions, the technical requirements for extracting, structuring and making sense of the data largely overlapped, and included: a) corpus-related document pre-processing and cleansing (e.g., removing misleading form headings from scanned documents); b) common medical terminology compilation and recognition (e.g., the antipsychotic medication identification problem is almost the same in [10] and [11]); and c) deriving patient-level clinical signals from document-level NLP annotations (e.g. understanding that a medication prescribed at admission time was actually removed from the patient’s discharge medication list).

As unstructured EHR data is inevitably needed by many research projects and clinical studies, more cost-effective and systematic solutions are needed to address the common challenges faced by different use cases, while also ensuring that study-specific requirements are not compromised by the unified approach.

To address such challenges, we propose SemEHR - a semantic search and analytical system generating a complete and processable view of a patient from his/her clinical notes.

- To realise a *general-purpose* biomedical information extraction (IE) system on EHRs, there are at least three fundamental challenges: a) syntactic heterogeneity: how to effectively access multi-modal/multisource EHR data that are almost certainly heterogeneous in formats, data models and access interfaces; b) knowledge coverage: how to cover all possible biomedical concepts that are required by potential use cases; c) context capturing: how to represent and capture the contexts associated with extracted concepts, and which are critical to understanding the clinical domain. To address these challenges, SemEHR architects a production infrastructure that integrates our previous work in the CogStack pipeline [14] to harmonise and cleanse heterogeneous records, using them to identify contextualised mentions (negation, temporality and experiencer) of a wide range of biomedical concepts including SNOMED CT^1^, ICD-10^2^, LOINC^3^ and Drug Ontology^4^. In addition, SemEHR automatically associates semantic types of annotations and their clinical contexts (derived from containing documents or sections) with dedicated extraction rules, which enables better IE capabilities such as populating the structured vital sign data from observation notes.
- It is well appreciated that one-size-fits-all approaches need to be adapted to work effectively in different scenarios. Therefore, to serve different use cases well, we require the capability to to extend the terminology of the general-purpose IE system to cover unseen concepts, deal with language specificities in a subcorpus, support use-case-specific extraction requirements and enable performance fine-tuning, e.g. by incorporating specific knowledge or researchers’ expertise. SemEHR provides a study-based (use-case-specific) learning engine enabling iterative learning and feedback. It collects user feedback and uses rule-based and machine learning techniques to tackle study-specific challenges and requirements in a continuous manner.
- A few hurdles prevent the effective consumption of extracted data from general-purpose IE systems in scientific research and clinical studies. To fulfil requirements from various studies, general-purpose IE systems are inevitable to adopt large terminologies, which users might/could not be familiar with. This poses challenges in a) mapping look-up concepts to terminology terms; b) translating clinical relations to term associations; c) exploiting terminology semantics for bring ‘unexpected/unperceived’ new insights. At the consumption level, SemEHR implements an ontology-based semantic search component to tackle such challenges.
- Lastly, and probably most importantly, EHRs represent a timeline of multiple patient interactions with services. As such, The document level IE results should be integrated at patient level to incorporate temporal and macro-contextual information (which reports, which visits, etc. as opposed to the sentence-based contextual information discussed above). Only after this integration is the EHR IE task complete. However, this requires a thorough understanding of the EHR system. SemEHR provides a multiperspective view for each patient by assembling NLP annotations at patient-level as longitudinal views and compiling structured medical profiles. Both NLP results and the patient timeline are made available via an ontology-based search system, which effectively turns common IE tasks into semantic search queries. The interface provides a multi-perspective view for each patient by assembling NLP annotations at patient-level as longitudinal views and compiling structured medical profiles.

## Method

### Data model and longitudinal patient views

As depicted in Figure 1a, SemEHR is built upon six types of entities - Patient, Clinical Note, Concept, Concept Mention, Medical Profile and Profile Aspect. Each *patient* is associated with a list of dated and typed *clinical notes*. From these notes, SemEHR identifies *mentions* of a wide range of biomedical *concepts* from the Unified Medical Language System (UMLS) [15,16], a compendium of many controlled vocabularies including SNOMED CT, ICD-10, LOINC, Drug Ontology and Gene Ontology. By analysing the context of its appearance, each *mention* is associated with three dimensional contextual information - negation, temporality and experiencer. Highlighted in green in Figure 1a, the associations between concepts (e.g. *Steatohepatitis is a liver disease; Ribavirin is a drug for treating Hepatitis C*) are made available for conducting semantically enriched computations by incorporating the various biomedical ontologies and Linked Open Data^5^ such as DBpedia [17] and Wikidata [18]. SemEHR derives periodical *medical profiles* from clinical notes belonging to a patient, - automatically generated medical summaries consisting of a set of *profile aspects* (sections describing different aspects of a medical profile, e.g. past medical history, medication and etc.) for a defined period of time. *Concept mentions* are assigned to these *aspects* according to their appearances in the original clinical notes. As the yellow boxes of Figure 1a show, SemEHR entities are mapped to Fast Healthcare Interoperability Resources (FHIR)^6^ entities whenever possible.

**Figure 1a.**
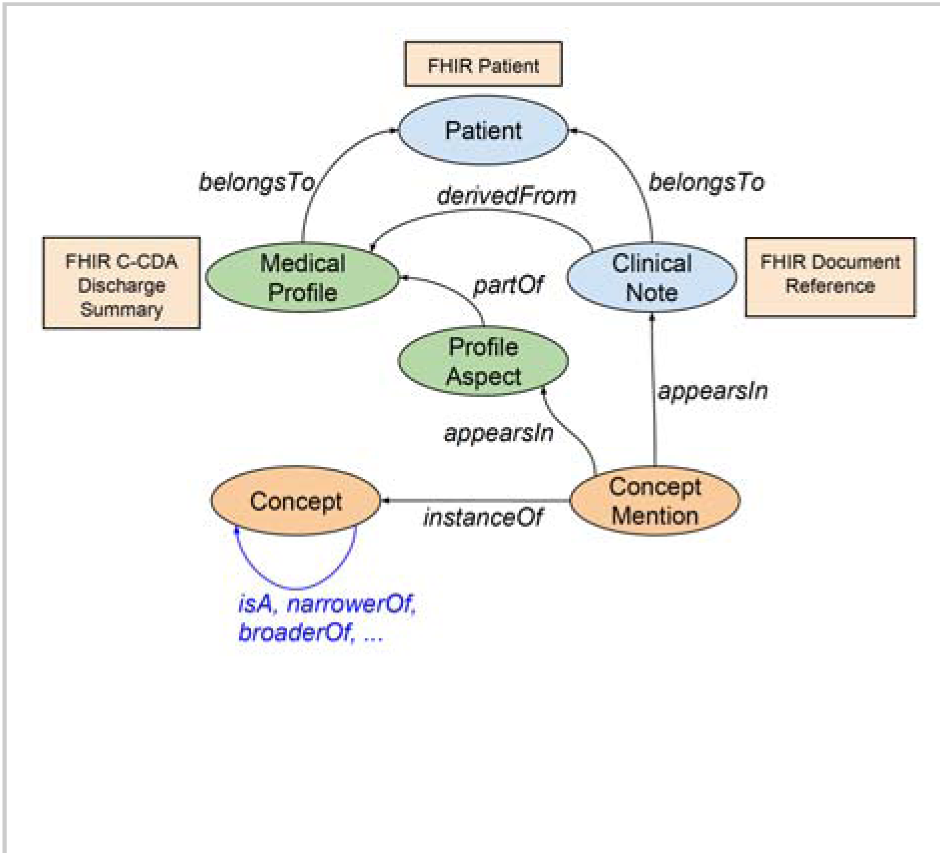
SemEHR Data Model: Entities (Patient, Clinical Note, Concept and Concept Mentions) and their associations.

Based on this data model, SemEHR populates two longitudinal views (shown in Figure 1b) for each patient. As shown in the upper part of Figure 1b, the first view is generated directly from the raw data. Concept mentions are organised in a list of clinical notes that are located on a timeline according to their date attributes (e.g. the created datetime of the clinical notes). Wherever possible, types of clinical notes (such as GP letter, Radiology or Discharge summary) are presented.

**Figure 1b.**
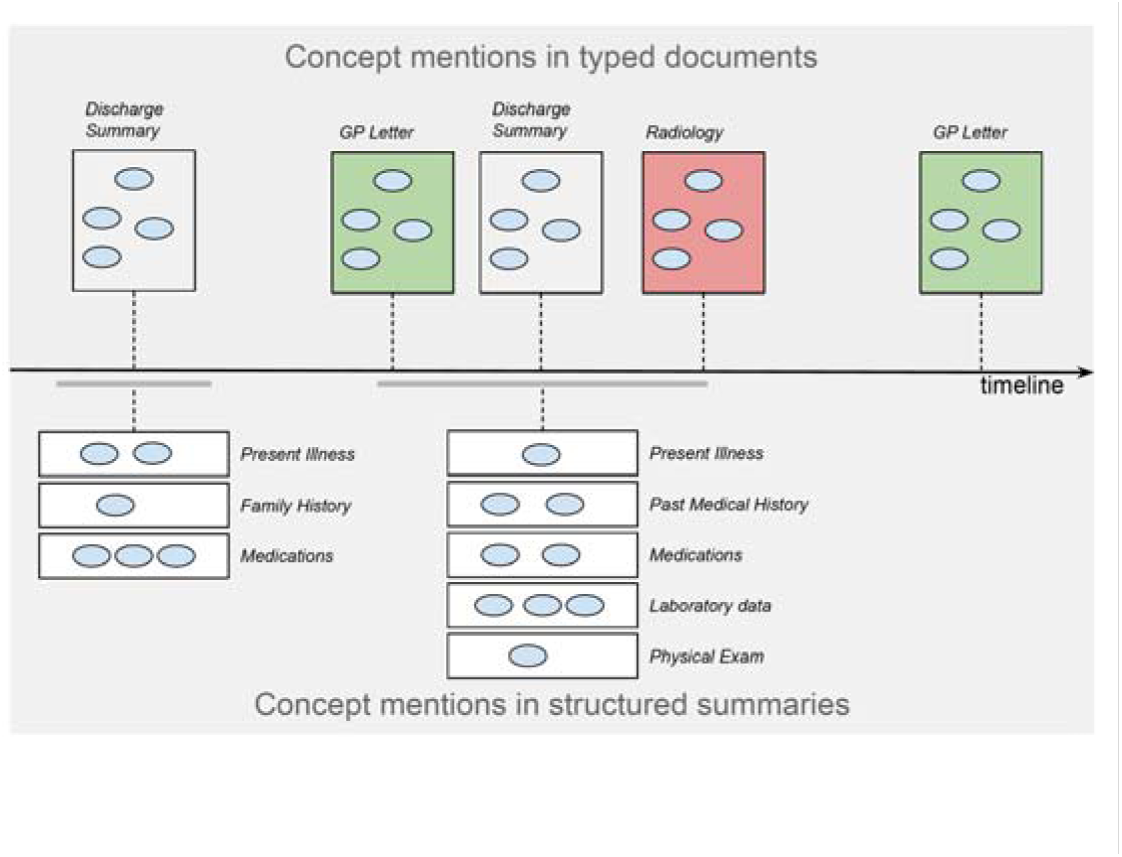
SemEHR generates two longitudinal views for each patient: concept mentions grouped in typed and dated documents (upper part); and concept mentions grouped in structured (discharge) summaries (lower part).

The second view (lower part of Figure 1b) is designed to convey structured summaries for a patient, each of which summarises the patient’s medical histories/conditions in a period of time (e.g. the hospital stay of an inpatient). A summary is composed of groups of concept mentions, where each group is about one particular aspect of the patient’s medical profile, e.g. past medical history, medications or physical exams. Preferably, such summaries are derived from discharge summaries. When discharge summaries are not available, an automated summarisation approach is applied to generate the summaries based on the concept types’ and concept mentions’ contextual information. Automated summaries are differentiated from those generated from discharge summaries. Supplementary material 2 describes the detailed process of automated medical profile generation.

#### Architecture: generic and adaptive information extraction and retrieval

As illustrated in Figure 2, SemEHR is composed of three subsystems -the producing subsystem, the continuous learning subsystem and the consuming subsystem.

**Figure 2.**
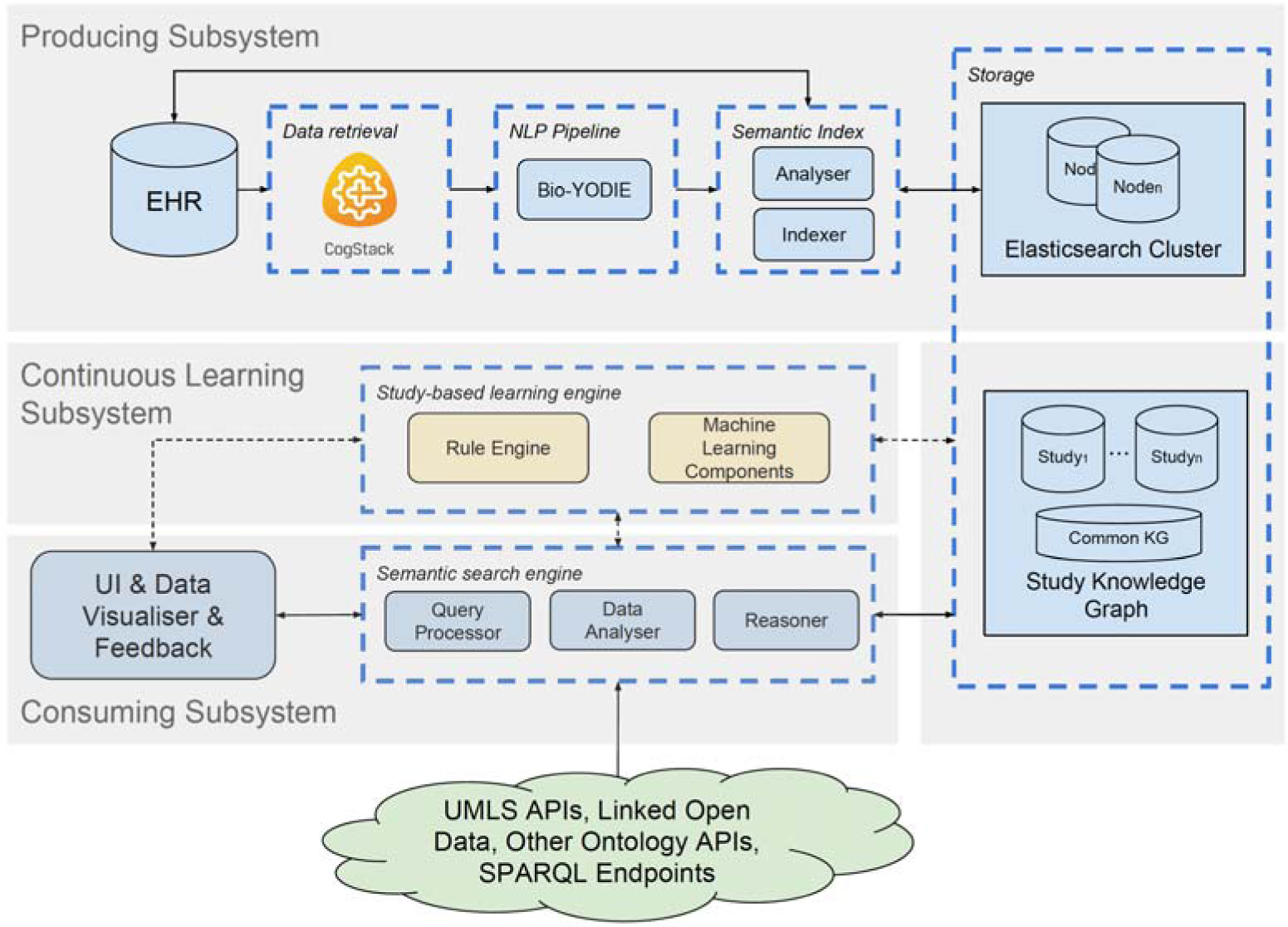
The architecture of SemEHR is composed of three subsystems: 1) the producing subsystem (upper part of the figure) - the creation of SemEHR semantic index by harmonising, natural language processing and indexing EHR data; 2) continuous learning subsystem - addressing study-specific requirements and supporting fine-tuning for separate studies 3) the consuming subsystem (lower part) - supporting tailored care, patient recruitment and clinical research by semantic searching and study-based continuous learning;

##### The Producing Subsystem

Essentially, the producing subsystem extracts free-text clinical notes from heterogenous underlying EHR systems, populating the data model described in the previous section. This task is performed in three main steps: data retrieval, information extraction and semantic indexing. CogStack[14], a data harmonisation and enterprise search toolkit for EHRs, is adopted in the data retrieval step to provide a unified interface to unstructured EHR data, which is often very heterogeneous in format and distributed in storage. Each document that flows out from the data retrieval component is fed into the NLP pipeline, which embeds Bio-YODIE^7^, an NLP pipeline dedicated to annotating UMLS concepts in clinical notes (‘documents’ for short hereafter). Emerging from the NLP pipeline are the documents and concept mentions extracted from them, both of which are then analysed by the Semantic Index component before being indexed. The analysis involves deriving document types (e.g. Radiology, GP letter and discharge summary), parsing document structure, e.g. identifying headed blocks from discharge summaries, and associating concept mentions with document structures. The analysis results, document content and NLP outputs are finally indexed by an Elasticsearch^7^ cluster. Patient-level summaries are generated as described in the previous section. These summaries are updated as new documents are added to the index.

SemEHR aims to produce annotations with accurate contextual information. Three components work collectively to achieve this goal - the Bio-YODIE pipeline captures the sentence/paragraph level contexts (e.g., negation, hypothetical mentions); the semantic index’s analyser brings in section/document level context (e.g., past medical history, laboratory results); the continuous learning subsystem (to be described in next subsection) learns the contexts from user assessed annotations (see Supplementary Material 1 for detail).

##### The Continuous Learning Subsystem

To accommodate the uniqueness of the IE requirements of different studies, SemEHR is designed with a continuous learning subsystem to iteratively address study-specific issues. The system collects and analyses user feedback from an annotation component embedded within the user interface. Based on the analysed feedback, two components are used to improve the IE results. The first is a rule engine, which generates and applies rules for filtering out unwanted results, e.g. removing concept mentions based on its original string or surrounding texts. The second component is a machine learning engine (a bidirectional recurrent neural network model), which takes user feedback as training data, applies the trained model on the study’s corpus and populates a confidence value for each concept mention. Confidence values are used as a quantified indicator in analytic components for populating results. The user interface for collecting feedback and continuous learning mechanisms are explained in detail in Supplementary Material 1.

**Figure 3.**
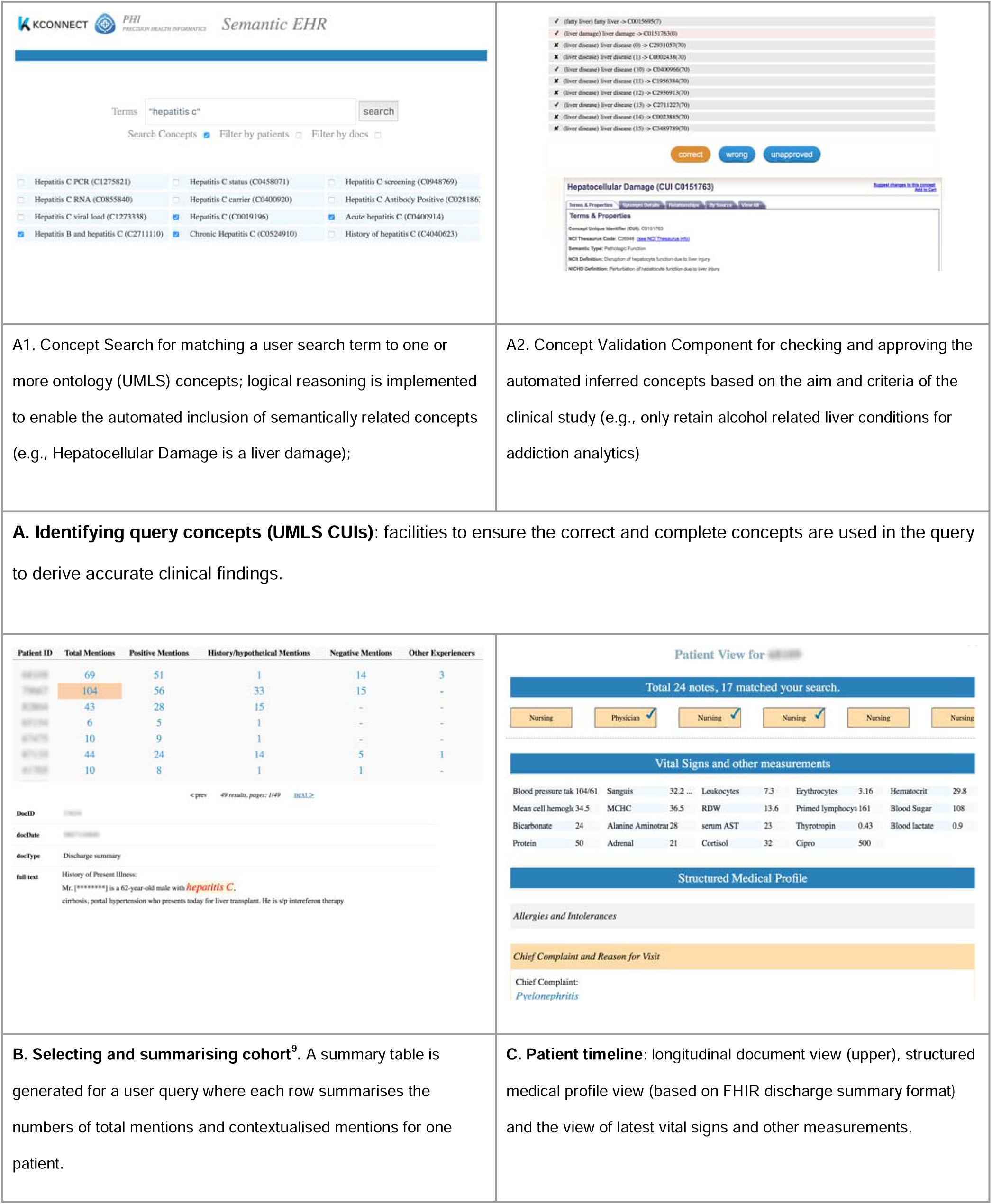
Screenshots of key functionalities provided by the consuming subsystem.

##### The Consuming Subsystem

This subsystem consists of a set of components that utilise information extraction results and clinical knowledge (accessed from biomedical ontology and Linked Open Data APIs) to support tasks such as patient characterisation or trial recruitment. A consuming task is called a ‘study’ in SemEHR. Each study will have its own storage within SemEHR’s Study Knowledge Graph (the bottom of Storage Section in Figure 2), which stores its study parameters (e.g., cohort definition and metadata), search settings (e.g., query concepts), study results (e.g., selected cohort and exported features) and customised rules (e.g., regular expressions to remove unwanted annotations). There is also a common knowledge graph (KG) (Common KG in Figure 2), where sharable knowledge or efforts (such as manually selected concepts of alcohol-related liver diseases or post processing rules for improving NLP results) are made available to other studies.

Key functionalities of the consuming subsystem include:

- *Translating search terms to query concepts.* This translates the user’s keyword searches (which are often ambiguous or incomplete) into semantically clear concepts (identified using UMLS CUIs). The correct translation is essential to ensure the soundness and completeness of search and analytics results. Unfortunately, in the clinical scenario, it is often not a trivial task to compile an accurate and complete concept list even just for a single clinical signal. For example, one SemEHR case study needs to look up patients with alcohol-related liver diseases. Given a general clinical term such as “liver disease”, it would be time consuming to compile a list of all subtypes of liver disease, which are also alcohol-related. As depicted in section A of Figure 3, SemEHR provides two functions for supporting concept translation: A1) matching search terms to concepts, which is enhanced with logical reasoning to automatically include semantically related concepts and EHR-based exclusion to remove concepts that do not exist in EHRs of the study cohort; A2) validating automatically populated lists, which is to allow manual assessment by the researchers.
- *Selecting and summarising a cohort.* Each query submitted to SemEHR will result in a cohort - a list of patients who match the query. As shown in section B of Figure 3, a summary table is generated for the matching cohort. Each row summarises a patient, and the first column shows the patient ID. The second one shows the total number of mentions of the search concepts within this patient’s EHRs, followed by numbers of four contextualised mentions - positive mentions, history/hypothetical mentions, negated mentions and mentions associated with other experiencers. Clicking on the numbers brings the user to the clinical notes where corresponding mentions are highlighted (lower part of Figure 3 section B).
- *Generating patient views and structured medical profile.* As a generic information extraction and retrieval platform, SemEHR processes all EHR records for a patient and tries to identify a wide range of biomedical concepts from them. This enables it to produce a panorama for each patient. As shown in section C of Figure 3, three different views are generated for each patient.
  - The first view is the longitudinal document view (the upper part of Figure 3 section C), which lists all patient documents in chronological order, labels documents using their types and ticks those documents that matched the query. This view delivers the abundance of a patient’s records, the prevalence of matched documents and their temporal distributions.
  - The second view is the structured medical profile (lower part of Figure 3 section C) that is automatically derived from patients’ clinical notes and structured using extended FHIR discharge summary format^10^. This structured summary enhances SemEHR’s searching and IE ability. For example, by constraining the search field to “Family History”, one can get a cohort with a family history of a certain disorder. In addition, knowing a piece of text appeared in the “Hospital Discharge Physical”, sophisticated rules can be applied to extract more structured data such as vital signs.
  - The third view is the view of vital signs and other measurements (middle part of Figure 3 section C). This is automatically generated by applying IE rules on the latest structured summary of a patient.

Based on these key functionalities, SemEHR provides a set of search interfaces to surface the clinical variables hidden in clinical notes. A typical query such as ‘return all patients with a family history of hepatitis c’, which previously might have required the end user to have NLP expertise, e.g., doing named entity recognition for ‘heptatitis c’ that must be mentioned in the context of ‘family history’. Using SemEHR, the end user can put in a simple keyword search: “hepatitis c”. To fulfill this search, SemEHR will pull out the cohort of relevant patients, populate patient-level summaries, i.e. numbers of contextualised concept mentions (e.g., 2nd patient has 16 total mentions of the disease, 15 of them were positive and 1 was about a family member), and provide a link to each mention in the original source clinical note (similar to the UI illustrated in Figure 3B).

## Results

### Studies conducted on CRIS data of South London and Maudsley Hospital

SemEHR has been deployed on the anonymised psychiatric records database CRIS [8], which contains a total of 18 million free-text documents from South London and Maudsley Hospital - one of Europe’s Largest Mental Health Provider (serving 1.2m residents). In the CRIS clinical notes, SemEHR identified 46 million mentions of concepts, the predominant ones being Pharmacologic Substances (16 million), Mental or Behavioural Dysfunction (12 million) and Sign or Symptom (3.8 million). In a CRIS-based liver disease study, SemEHR identified 94 instances out of 100 Hepatitis C positive patients who were manually annotated (based on structured blood test data). In an HIV study, a random 1,000 patient cohort was selected, and SemEHR identified 21 out of 23 true positive (verifiable via structured blood test data) HIV patients using two search concepts - HIV Pos (UMLS code: C0019699) (20 TP) and HIV diagnosis (UMLS code: C0920550) (8 TP). SemEHR integrates document-level NLP annotations to the patient level for generating an integral view for patients. Table 1 presents the results of two experiments designed to evaluate the effectiveness of such integration on two case studies - Hepatitis C and HIV. The results show that the number of positive mentions of diseases at patient level is a good feature for supervised learning methods (Naive Bayes or decision table) to classify whether a patient suffers from a disease or not.

**Table 1.**
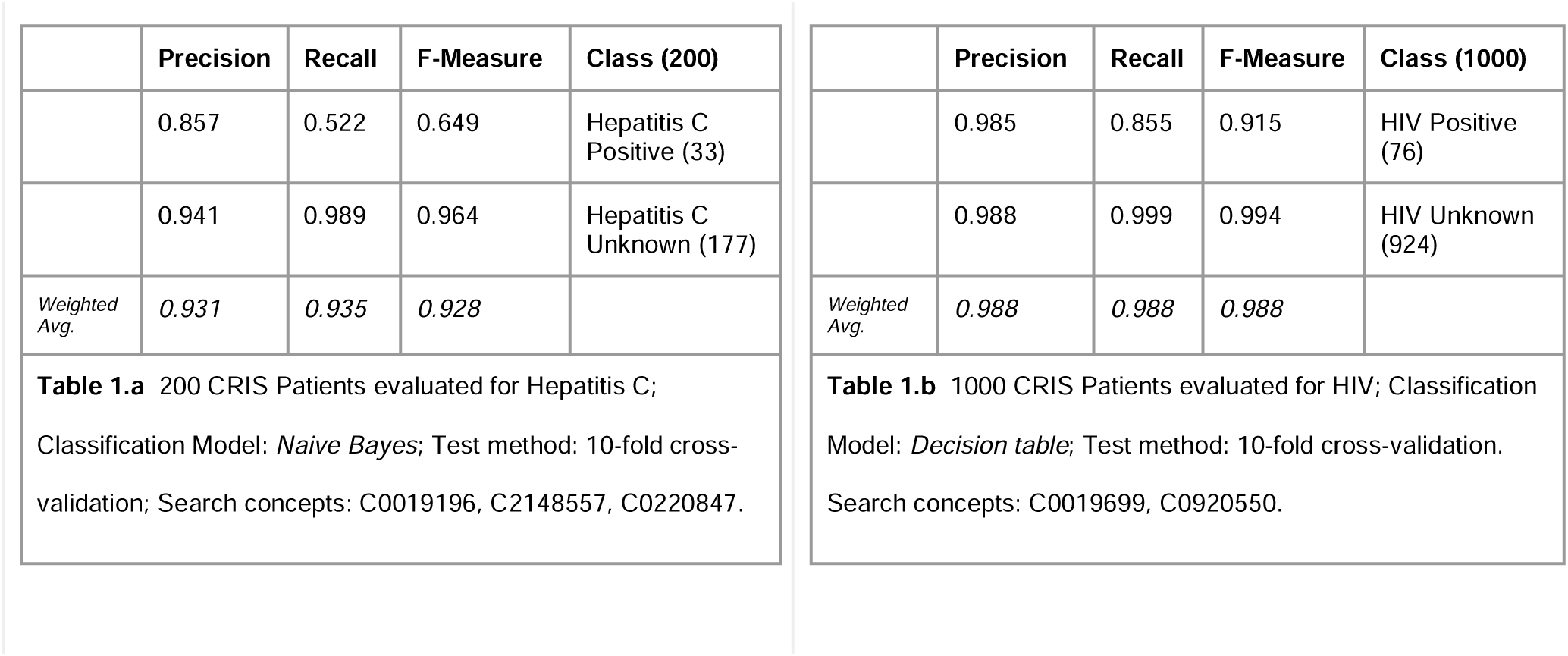
Given a disease (identified by one or more UMLS concepts, i.e. search concepts), SemEHR can generate a summary table for a cohort of patients, which, for each patient, gives the number of positive mentions of the search concepts within all his/her EHR documents. Using this number as the only feature, we classify whether a patient suffers from a disease or not. Table 1.a presents the result of a 200-patient-cohort for hepatitis C infection and Table 1.b shows the result of a 1000-patient-cohort for tHIV.

### Study conducted at King’s College Hospital, London

At King’s College Hospital, SemEHR is being used to assess the eligibility of, and subsequently recruit patients into the 100,000 Genomics England Project^11^. Here, an open SPARQL endpoint is integrated to map UMLS concepts to Human Phenotype Ontology (HPO) terms, inclusion criteria for recruitment and the concepts necessary to populate complex phenotype models. The preliminary validation study suggests that the tool is able to validate previously submitted cases and is very fast (providing results within seconds) in searching phenotypes; an operation that previously required manual assessments through patient records. For example, the time for recruitment criteria checking of a patient is reduced significantly from days to minutes for dermatology disorders, of which the inclusion/exclusion criteria contain 120 phenotypes in average. In addition, the semantic reasoning (e.g., expanding searching concepts with more specific concepts) has been found to be helpful for identifying two specific phenotypes, namely Neutropenia and Hypertension.

### Studies conducted on MIMIC-III data

We deployed a SemEHR instance on MIMIC-III [19], an intensive care EHR dataset anonymised from two US-based hospitals and made public for research purposes. MIMIC-III contains about 2 million free-text clinical notes and comprises very good structured data including high-resolution laboratory measurements for most patients. To evaluate the performance of SemEHR’s structured medical profile, we randomly selected 100 patients and assessed the accuracy of automatically extracted laboratory measurements in their SemEHR medical profiles. The results are presented in Table 2. Eleven types of laboratory measurements were manually selected for this evaluation, which contains popular tests such as Hematocrit and relatively rare ones such as blood urea. First, we compared the extracted measurement values with those stored in the MIMIC-III structured data. A patient usually has multiple values of the same measurement that were measured/tested at different times and it should be noted that as long as the extracted value appears within the list of all values from the structured data, the extraction is deemed correct; otherwise incorrect. The result of the first step is presented in the second last column. The average accuracy using structured data verification is 89%. For those incorrect extractions, we applied a second step of manual assessment. This step identified some false negative results from the first step caused by reasons including decimal rounding (3 cases), different units (2 cases) and missing laboratory events in structured data (6 cases). The accuracies based on the manual verification are reported in the last column of Table 2. The average accuracy was improved to 97%.

**Table 2.**
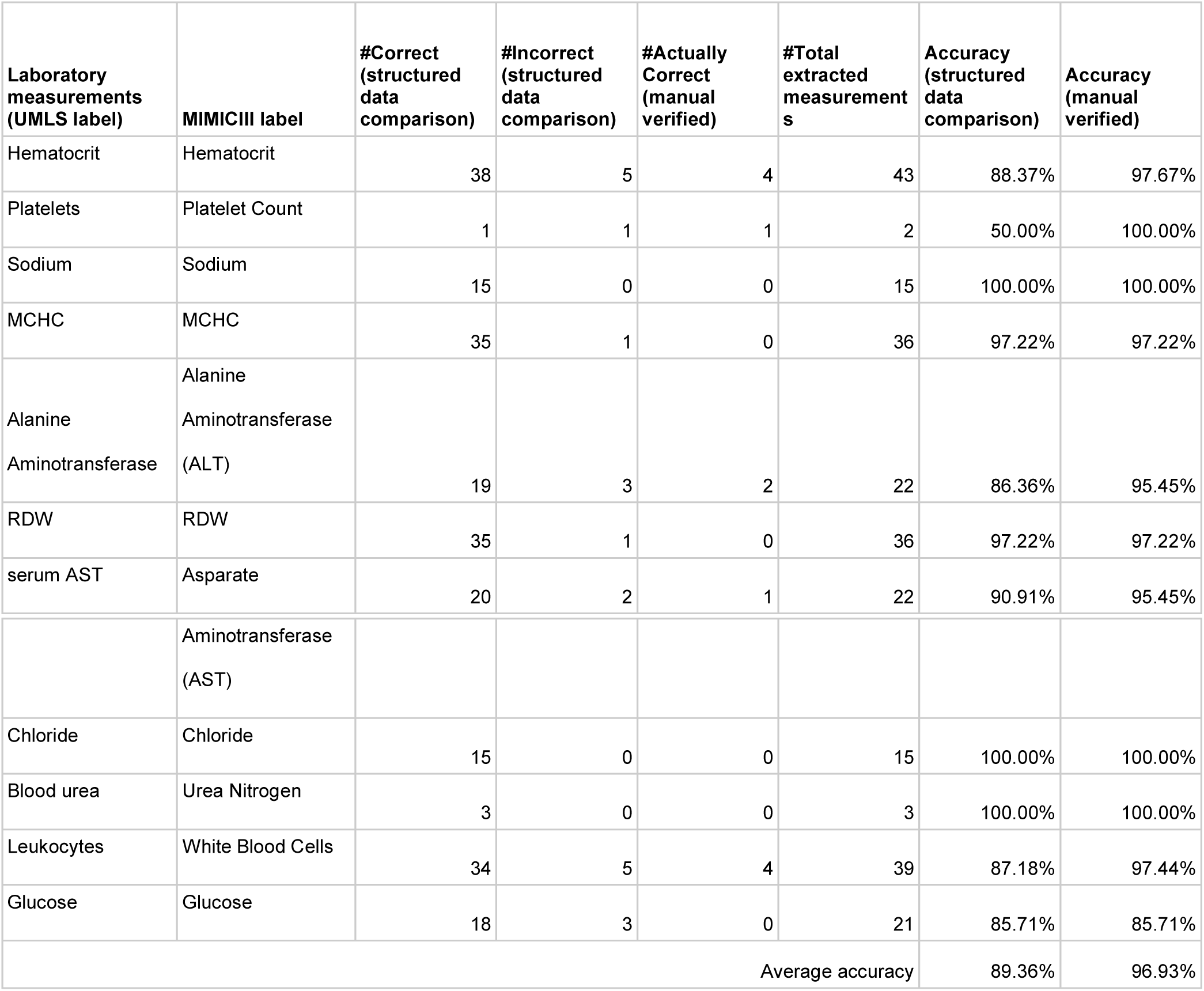
The performance of SemEHR laboratory measurement extraction on MIMICIII data: 11 measurements are studied (first column); 100 patients were randomly selected for this study; the extracted results were assessed by two steps: a) comparing with the structured data (querying labevents table in MIMICIII) - accuracy reported in the 7th column; and b) manual checking on not-matched items in the first step (accuracy reported in the last column).

The manual verification revealed that extracting vital signs from clinical notes can complement structured data in MIMIC-III - there are 6 cases where the measurements are extracted from free texts but missing in the structured data. In general, SemEHR can reveal various types of structured data that are usually not or cannot be recorded in structured EHRs, such as family history and social history. In table 3, for the above 100 randomly-selected patients, we summarise the number of semantic entities identified in 5 sections of SemEHR medical profiles which are usually not recorded in structured EHRs.

**Table 3.** The number of extracted semantic entities in 5 sections of SemEHR medical profiles of the 100 randomly selected MIMIC-III patients, which are usually not recorded in structured EHRs.

|  | Admission Medications |  | Family History |  | Social History |  | History of Past Illness |  | Hospital Discharge Instructions |  |
| --- | --- | --- | --- | --- | --- | --- | --- | --- | --- | --- |
| #Total annotations | 1,475 |  | 156 |  | 445 |  | 1,575 |  | 1,162 |  |
| Top 5 semantic types by frequency | Temporal Concept | 442 | Finding | 42 | Finding | 132 | Disease or Syndrome | 337 | Clinical Attribute | 359 |
|  | Pharmacologic Substance | 393 | Disease or Syndrome | 33 | Temporal Concept | 86 | Finding | 189 | Temporal Concept | 158 |
|  | Finding | 194 | Neoplastic Process | 28 | Pharmacologic Substance | 58 | Temporal Concept | 182 | Health Care Related Organization | 133 |
|  | Clinical Drug | 121 | Pharmacologic Substance | 14 | Clinical Attribute | 30 | Therapeutic or Preventive Procedure | 180 | Health Care Activity | 126 |
|  | Health Care Related Organization | 51 | Clinical Attribute | 9 | Individual Behavior | 28 | Body Part, Organ, or Organ Component | 96 | Finding | 79 |

## Discussion

SemEHR has been deployed or in the process of deploying to a number of NHS Trusts’ EHR systems including South London and Maudsley, King’s College Hospital, University College London Hospitals and Guy’s Hospital. Results and feedback from the multiple SemEHR use cases have shown its effectiveness in automating lengthy manual tasks without jeopardising accuracy. Queries are returned at a rate rapid enough to enable iterative tailoring to achieve high specificity. Moreover, according to our case studies at SLaM, SemEHR has achieved similar accuracy to bespoke NLP applications built upon TextHunter [13]. Powered by ontological semantics, researchers can make use of semantically associated concepts to improve results, e.g. in the CRIS-based liver disease study, the inclusion of 8 drugs used for treating liver diseases have helped to find more patients.

Our case studies have shown that building a unified framework like SemEHR realises a more cost-effective approach to dealing with common information extraction challenges and significantly lowers the barrier for researchers, coders and clinicians to access knowledge residing within unstructured clinical notes. SemEHR has great potential in enabling the efficient and effective secondary use of EHRs to improve healthcare services. Furthermore, SemEHR-like systems initiate a collaborative learning platform as advocated by Moseley and et al. [20], enabling studies to be conducted in a cooperative way rather than resources remaining in isolated silos.

SemEHR provides different patient views, with the aim of presenting a more continuous representation of the patient’s treatment timeline. Such views may reveal data quality issues [21,22] to researchers or clinicians so that necessary actions can be taken before deriving conclusions. For example, the longitudinal document view gives a quick overview of how abundant or detailed a patient’s EHR is, which helps identify patients who have incomplete records and might need to be removed from studies. However, data quality issues such as data incompleteness, inconsistency and inaccuracy need to be addressed in a systematic way; making users aware of the potential issues is only the first step. In our future work, we will investigate approaches to tackle challenges such as automated patient-level consistency checking, bearing in mind that some of the challenges require wider-scope, e.g. institution-level, attention [23,24].

## Conclusion

In this paper, we presented SemEHR, a unified information extraction (IE) and semantic search system for obtaining clinical insight from unstructured clinical notes. With a dedicated architecture and the incorporation of semantic analytics, SemEHR effectively turns IE tasks into (iterative) ontology-based searches, which significantly lowers the barriers to secondary use of unstructured EHR data. The system has been deployed in several NHS hospitals in the UK and a number of case studies have been initiated, including patient recruitment for the UK government’s 100,000 Genomics England project. Results and feedback demonstrate that SemEHR can efficiently perform the task of cohort selection and patient characterisation with high accuracy. SemEHR is open source; all non-sensitive data relating to its verifications has been published in its online repository: https://github.com/CogStack/SemEHR.

## Contributorship Statement

HW, IK, AF, AR, GG, RJ, ZI, and AA were involved in development of SemEHR or components that were used by the system. GT and KIM led the liver disease and HIV studies on South London and Maudsley (SLaM) EHR. RS led an autoimmune study on SLaM EHR; CS and DG led the 100,000 Genomes Project study at King’s College Hospital. MB and RS provided the access and computational resources for accessing SLaM EHR. AR, RS, and RJBD secured funding for this research. All authors contributed to the abstract.

## Footnotes

1 http://www.snomed.org/snomed-ct

2 http://apps.who.int/classifications/icd10/browse/2010/en

3 https://loinc.org/

4 https://ontology.atlassian.net/wiki/spaces/DRON/overview

5 https://en.wikipedia.org/wiki/Linked_data

6 https://www.hl7.org/fhir/overview.html

7 https://gate.ac.uk/applications/bio-yodie.html, developed as part of EU KConnect project in which GG, AR, HW, RS, RD are involved.

8 https://www.elastic.co/products/elasticsearch

9 The full text in the screenshot has been deliberately rewritten to avoid leaking sensitive patient data.

10 23 sections of FHIR discharge summary (http://hl7.org/fhir/us/ccda/2017Jan/StructureDefinition-CCDA-on-FHIR-Discharge-Summary.html) are extended with additional 8 headings.

11 https://www.genomicsengland.co.uk/

